# Temporal Dynamics and Performance Association of the *Tetrasphaera*-Enriched Microbiome for Enhanced Biological Phosphorus Removal

**DOI:** 10.1101/2022.08.23.504879

**Authors:** Hui Wang, Yubo Wang, Guoqing Zhang, Ze Zhao, Feng Ju

**Affiliations:** Environmental Science and Engineering Department, Zhejiang University, Hangzhou 310012, Zhejiang Province, China; Key Laboratory of Coastal Environment and Resources of Zhejiang Province, School of Engineering, Westlake University, 18 Shilongshan Road, Hangzhou 310024, Zhejiang Province, China; Institute of Advanced Technology, Westlake Institute for Advanced Study, 18 Shilongshan Road, Hangzhou 310024, China; Center of Synthetic Biology and Integrated Bioengineering, School of Engineering, Westlake University, Hangzhou 310024 Zhejiang Province, China

**Keywords:** Enhanced biological phosphorus removal (EBPR), Polyphosphate-accumulating organisms (PAOs), *Tetrasphaera*, Microbiome, Phosphorus recovery

## Abstract

*Tetrasphaera* were recently identified based on the 16S rRNA gene as among the most abundant polyphosphate-accumulating organisms (PAOs) in global full-scale wastewater treatment plants (WWTPs) with enhanced biological phosphorus removal (EBPR). However, it is unclear how *Tetrasphaera* PAOs are selectively enriched in the context of the EBPR microbiome. In this study, an EBPR microbiome enriched with *Tetrasphaera* (accounting for 40% of 16S sequences on day 113) was built using a top-down design approach featuring multicarbon sources and a low dosage of allylthiourea. The microbiome showed enhanced nutrient removal (P removal ~85% and N removal ~80%) and increased P recovery (up to 23.2 times) compared with the seeding activated sludge from a local full-scale WWTP. The supply of 1 mg/L allylthiourea promoted the coselection of *Tetrasphaera* PAOs and *Microlunatus* PAOs and sharply reduced the relative abundance of both ammonia oxidizer *Nitrosomonas* and putative competitors *Brevundimonas* and *Paracoccus*, facilitating the establishment of the EBPR microbiome. Based on 16S rRNA gene analysis, a putative novel PAO species, EBPR-ASV0001, was identified with *Tetrasphaera japonica* as its closest relative. This study provides new knowledge on the establishment of a *Tetrasphaera*-enriched microbiome facilitated by allylthiourea, which can be further exploited to guide future process upgrading and optimization to achieve and/or enhance simultaneous biological phosphorus and nitrogen removal from high-concentration wastewater.

## 1. Introduction

Phosphorus (P) eutrophication and P mineral resource depletion are increasingly serious global problems. Wastewater treatment plants (WWTPs) are both the sink and source for P [1, 2]. The bioprocess known as enhanced biological phosphorus removal (EBPR) can effectively promote P removal and recovery in WWTPs. It relies on a specific microbial functional guild called polyphosphate-accumulating organisms (PAOs) [3, 4]. Until now, most studies of EBPR microbiology have focused on *Ca.* Accumulibacter PAOs, which are commonly found in full-scale WWTPs and are widely enriched in lab-scale reactors using volatile fatty acids as carbon sources [4]. Because of the complex wastewater compositions and luxuriant growth conditions, full-scale WWTPs can provide diverse nutritional niches to accommodate various functional guilds [5, 6]. PAOs other than *Ca.* Accumulibacter (e.g., *Tetrasphaera* and *Dechloromonas*) are increasingly found to play an important but previously underappreciated role in full-scale systems [4, 7, 8], which encourages continued and follow-up efforts to explore the emerging aspects of the microbiology and ecology of the EBPR process.

*Tetrasphaera* PAOs have gained increasing attention due to their prevalence in global WWTPs as both important contributors to P removal and potential reservoirs for P recovery [4, 7]. For example, *Tetrasphaera* were found to be the most abundant PAOs in 18 Danish full-scale EBPR WWTPs [7] and were further revealed by Raman-FISH analysis to have a comparable or even higher P removal contribution than *Ca.* Accumulibacter PAOs [9]. In addition, *Tetrasphaera* PAOs are implicated in metabolizing various carbon sources (e.g., glucose and amino acids), which may favor their growth in WWTPs with fluctuating environmental conditions [10]. Although *Tetrasphaera* PAOs play an important role in the EBPR of full-scale WWTPs, how to selectively enrich and regulate them and how to enhance biological P removal (and thus recovery) remain elusive questions. Currently, *Tetrasphaera* PAOs, unlike the most intensively investigated *Ca.* Accumulibacter, are still at an infancy stage of research. Our knowledge of the phylogeny, ecophysiology, and metabolism of *Tetrasphaera* PAOs, especially their capacity for wastewater P uptake and removal, is limited [4, 9, 10]. The successful isolation of six *Tetrasphaera* strains partially filled these knowledge gaps, but the physiologies and metabolic characteristics of these isolates in pure culture cannot explain the interactive behaviors of *Tetrasphaera* PAOs in WWTPs [11]. In addition, the high diversity and abundance of *Tetrasphaera* PAOs in EBPR plants generally imply the existence of yet-to-be-discovered PAOs. Therefore, it is essential to explore the enrichment method and microbial interactions of *Tetrasphaera* PAOs in the context of the EBPR microbiome.

Until now, only a few studies have reported *Tetrasphaera*-enriched microbiomes that were fed amino acids as a sole carbon source [10, 12–14]. The enrichment strategy, process, and key biotic and abiotic factors are unclear, hindering the in-depth study of the ecophysiology and metabolism of *Tetrasphaera* PAOs. Notably, allylthiourea (C_4_H_8_N_2_S), a specific nitrification inhibitor, is empirically supplemented to facilitate the enrichment of model PAOs (mainly *Ca*. Accumulibacter) in lab-scale EBPR reactors by suppressing nitrite production by ammonia-oxidizing bacteria (AOB) to alleviate adverse competition from denitrifying bacteria [15, 16]. Because *Tetrasphaera* PAOs possess versatile metabolism, including the denitrifying capability, the effect of allylthiourea on enriching *Tetrasphaera* PAOs needs to be explored. Recently, a top-down approach has been proposed that uses intricately designed operation conditions (e.g., substrate composition, loading rate, and redox conditions) to manipulate an existing microbiome through ecological selection to perform the desired biological processes [17]. While this new methodological framework provides an ideal theoretical guide for the establishment of the *Tetrasphaera*-enriched microbiome, a practical strategy is lacking and needs to be established. Additionally, understanding of the microbiome dynamics and treatment performance during the *Tetrasphaera*-enriched microbiome establishment process is lacking, especially regarding the influence of biotic and abiotic factors and the link to EBPR performance (i.e., P removal and recovery). This knowledge is the key to the reproducible establishment and precise regulation of a *Tetrasphaera*-enriched microbiome to improve EBPR performance.

Given the abovementioned knowledge gaps, our study aims to i) establish and explore a top-down strategy for *Tetrasphaera*-enriched microbiome establishment and characterize its P removal and recovery potentials, ii) investigate the microbiome temporal dynamics and interaction patterns of the *Tetrasphaera*-enriched microbiome establishment process, and iii) quantify the contribution of biotic and abiotic factors to microbiome dynamics and treatment performance. In addition, the nexus between microbial composition, environmental (e.g., operational) conditions, and treatment performance of the EBPR microbiome as well as the effect of allylthiourea on the enrichment of *Tetrasphaera* PAOs and EBPR performance have been investigated. To the best of our knowledge, this is the first proof-of-concept trial study to investigate the approaches and mechanisms of the establishment of the *Tetrasphaera*-enriched microbiome from its strategy design to the potential evaluation of P recovery.

## 2. Materials and Methods

### 2.1. Bioreactor setup and operation

To enrich *Tetrasphaera* PAOs, a lab-scale sequencing batch reactor with a working volume of 10 L was established and inoculated with activated sludge collected from a local municipal WWTP in Hangzhou, China. The WWTP employs the A/A/O process with AlCl_3_ coagulating sedimentation in the primary setting tank, treating 50000 m^3^/d of domestic wastewater. The 8-h cycle of reactor operation consisted of four phases: 150 min of anaerobic treatment, 320 min of aerobic treatment, 5 min of settling, and 5 min of water discharge and feeding. Operation details are provided in the Supplementary Materials (Text S1).

### 2.2. Synthetic wastewater preparation and enrichment stages

The synthetic wastewater was prepared by mixing different volumes of stock solutions (A, B, C, and trace element solution, Table S1) to maintain a TOC concentration of 394.8 ± 34.7 mg/L, PO_4_^3−^-P concentration of 24.8 ± 3.9 mg/L, and NH_4_^+^-N concentration of 113.6 ± 36.9 mg/L. The characteristics of the prepared synthetic wastewater are similar to those of high-concentration wastewater (e.g., livestock wastewater [18]). Specifically, solution A (multicarbon sources) contained amicase (CAS No. 65072-00-6, Sigma‒Aldrich), glucose (CAS No. 50-99-7, HuShi), and sodium acetate (CAS No. 127-09-3, HuShi). Solution B (P sources) contained K_2_HPO_4_ and KH_2_PO_4_. Solution C (mineral medium) contained NH_4_Cl, MgSO_4_·7H_2_O, and CaCl_2_·2H_2_O. The trace element solution was prepared according to a previous study ^15^ and added to synthetic wastewater (1 mg/L). To investigate the effect of allylthiourea on *Tetrasphaera* PAOs enrichment and EBPR function, the entire reactor operation was divided into the following four stages with different dosages of allylthiourea:

- Stage I (day 0-73): In the natural enrichment phase, no allylthiourea (0 mg/L) was supplemented as a nitrification inhibitor.
- Stage II (day 74-120): In the reinforcement enrichment phase, allylthiourea was supplied at 1 mg/L to enhance the *Tetrasphaera*-related PAOs enrichment by partially inhibiting nitrification activity.
- Stage III (day 121-127): In the transient shock phase, the allylthiourea was added to 5 mg/L to strongly inhibit or suppress nitrification activity.
- Stage IV (day 128-170): In the recovery phase, the allylthiourea concentration was adjusted back to 1 mg/L.

### 2.3. Batch experiments

Three batch tests were performed following experimental conditions similar to the mother reactor. Each test was seeded with the biomass collected from the EBPR reactor on day 143 when the relative abundance of *Tetrasphaera* sequences reached 30.3%. The composition of synthetic wastewater was consistent with Section 2.2 but only amended with three dosage levels of allylthiourea: 0 mg/L, 1 mg/L, and 5 mg/L. The pH and DO concentration were monitored using a portable multiparameter analyzer (HQ40d, Hach, USA). Seven samples were taken at three time points of the anaerobic phase (13, 45, and 135 min) and four time points of the aerobic phase (30, 90, 210, and 320 min). The sludge samples were centrifuged at 3000 rpm for 3 mins, and then the supernatant was collected for further physicochemical analysis.

### 2.4. Chemical analysis

The collected water samples were pretreated using 0.22 μm Millex GP syringe-driven filters to eliminate the potential solids before analysis. PO_4_^3−^-P, TN, and NH_4_^+^-N were analyzed using a segmented flow analyzer (San^++^, SKALAR, Netherlands). The TOC was assessed by a TOC-L total organic carbon analyzer (Shimadzu, Japan). TS and VS were analyzed according to standard methods [19]. To evaluate the P recovery potential of the *Tetrasphaera*-enriched microbiome, the P weight percentage content in sludge ash was assessed and compared with the WWTP. Briefly, the sludge samples obtained from the *Tetrasphaera*-enriched reactor and local WWTP (the sampling site was the same as seeding sludge) were adjusted to the same TS content using pure water. Samples with a 50 mL volume were taken and put into a crucible for incineration (600 ℃, 8 h). Then, the ash was collected and mixed using a mortar. The P weight percentage content in the sludge ash was assessed by a scanning electron microscope equipped with an energy dispersive spectrometer (SEM‒EDS) (Hitachi, Japan). Every experiment was conducted with the triple treatments.

### 2.5. FISH analysis of polyphosphate-accumulating organisms

Fluorescence *in situ* hybridization (FISH) analysis was performed to investigate the spatial organization of targeted PAOs in microbiome samples according to a previous protocol with some modifications [20]. The detailed experimental flow and the information of the FISH probes can be found in the Supplementary Materials (Text S2 and Table S2). FISH samples were observed using a laser scanning confocal microscope (Zeiss, Zeiss LSM800, Germany).

### 2.6. DNA extraction, PCR amplification, and sequencing

Sludge samples were collected regularly at the end of the aerobic phase for DNA extraction. A total of 16 samples were collected over 170 days. DNA extraction was performed in duplicate from 1.5 ml collected reactor biomass using the Fast DNA SPIN Kit for Soil (MP Biomedicals). The duplicate DNA extracts were mixed equally for bacterial 16S rRNA gene high-throughput sequencing targeting the V1-V3 hypervariable regions (27 F: AGAGTTTGATCCTGGCTCAG and 534 R: ATTACCGCGGCTGCTGG) [21] using Illumina’s MiSeq platform with a paired-end (2 × 300) sequencing strategy at Guangdong Magigene Biotechnology Co., Ltd., China. The thermocycler (Bio-Rad S1000, Bio-Rad Laboratory) settings for PCR were initial denaturation at 95 °C for 2 min, 30 cycles of 95 °C for 20 s, 56 °C for 30 s, and 72 °C for 60 s, and final elongation at 72 °C for 5 min [21].

### 2.7. Bioinformatics analysis

Fastp was used to check the quality of the raw data with the sliding window parameter of −W 4 −M 20. The primers were removed by using cutadapt software according to the primer information at the beginning and end of the sequence [22]. The obtained clean data were merged using QIIME2 (v. 2020.6) pipeline [23] with the DADA2 algorithm [24]. Briefly, the following steps were performed: i) filter out noisy sequences, ii) correct errors in marginal sequences, iii) remove chimeras sequence and singletons, iv) join denoised paired-end reads, and v) conduct sequence dereplication using a command of qiime dada2 denoise-paired with the parameter of --p-trunc-len-f 250 and --p-trunc-len-r 247 to maintain at least 12 bp overlap. The amplicon sequence variant (ASV) table was generated based on 100% sequence similarity for 16S rRNA genes, which contained 1389 ASVs in total. To exclude spurious sequences from the downstream analysis, the rare feature (i.e., ASV) with a total sequence count of less than 3 was removed using a command of qiime feature-table filter-features with the parameter of --p-min-frequency 3. To determine if the richness of the samples was fully sequenced, alpha rarefaction was analyzed using a command of qiime diversity alpha-rarefaction and visualized by the qiime 2 view website (https://view.qiime2.org/). A specific taxonomic classifier was trained based on the Silva_138.1_SSURef_NR99 database [25] and used for taxonomic analysis.

### 2.8. Phylogenetic analysis

The most abundant ASVs assigned to *Tetrasphaera* were selected to construct the phylogenetic tree using MEGA X [26]. Multiple sequence alignment was conducted by MUSCLE [27] with the default parameter. The evolutionary history was inferred by the maximum likelihood method and Kimura 2-parameter model. Two *Nitrosomonas europaea* sequences were used as the outgroup.

### 2.9. Statistical and network analysis

The statistical analyses, including redundancy analysis (RDA), principal coordinate analysis (PCoA), Mantel test and PERMANOVA test, were performed using the R package vegan [28]. Co-occurrence networks were constructed using the ‘Co-occurrence_network.R’ script of package MbioAssy1.0 (https://github.com/emblab-westlake/MbioAssy1.0) [6, 29]. Detailed information on the co-occurrence network analysis is provided in the Supplementary Materials (Text S3). The network was visualized in Gephi 0.9.2 (https://gephi.org/).

## 3. Results and Discussion

### 3.1. The establishment strategy and performance of the *Tetrasphaera*-enriched microbiome

In this study, we established a strategy including four stages to selectively enrich *Tetrasphaera* PAOs based on the top-down design approach [17]. In stage I, the multicarbon sources (containing amicase, glucose, and sodium acetate) and periodic operational conditions (anaerobic/aerobic alternations) were designed as the selective tactics to conduct the initial enrichment. In stage II, 1 mg/L allylthiourea was introduced to facilitate enrichment by suppressing nitrification. In the following stages (stage III and IV), a transient shock with a high dosage of allylthiourea (5 mg/L) was applied to investigate the effect of allylthiourea dosage on the *Tetrasphaera*-enriched microbiome.

The treatment performance of nutrients (i.e., NH_4_^+^-N, TN, and PO_4_^3−^-P) and organic carbon (i.e., TOC) in wastewater followed over nearly six months (Fig. 1). In stage I, the reactor showed an overall decline in P removal efficiency (average 35%), but NH_4_^+^-N removal increased with prolonged operation (the highest NH_4_^+^-N removal was up to 92% on day 49). With the addition of allylthiourea (stage II), the efficiency of P removal significantly increased from 27.6% (day 73) to 86.2% (day 94) and then remained stable between 83.3% and 87.4%. The NH_4_^+^-N removal decreased significantly during day 73-90 (the lowest percentage was 3.9% on day 87) and then recovered during day 105-121 (the average removal percentage was 97.6±3.0%). When the concentration of allylthiourea was increased to 5 mg/L in stage III (transient shock to suppress nitrification, Fig. 1C), the efficiency of P removal presented a slightly decreasing trend (from 86.2% of day 122 to 73% of day 125), accompanying an obvious decrease and final failure in NH_4_^+^-N removal. With the downregulation of the allylthiourea level back to 1 mg/L on day 127 (stage IV), the efficiency of P removal was recovered to the original level (67.4% to 89.3%), and the efficiency of TN removal was stabilized at approximately 80% on day 148-170. In addition, the utilization and degradation of organic carbon during the 170-day operation remained stable, and approximately 95% of the carbon source could be absorbed and utilized (Fig. 1B).

**Fig. 1.**
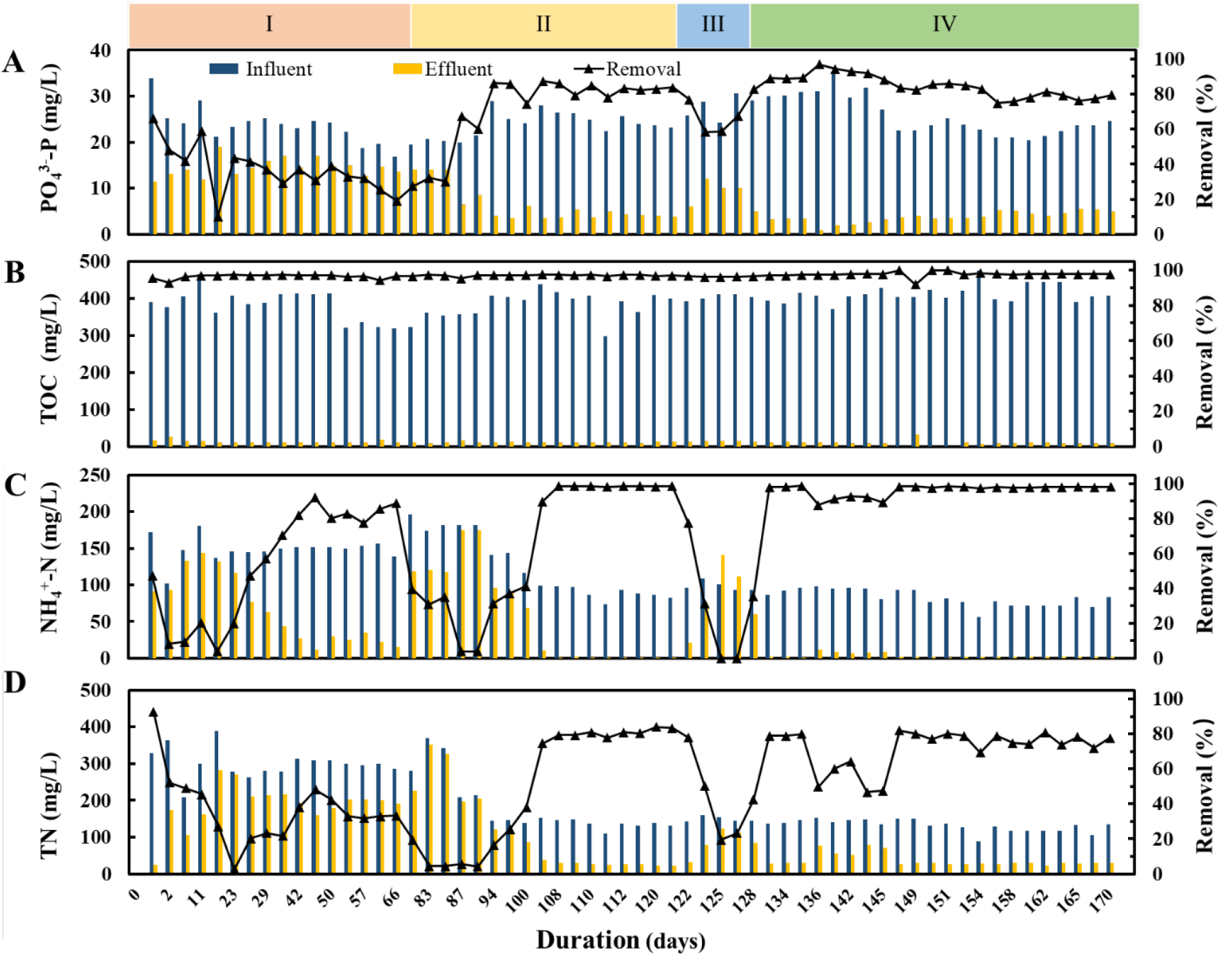
The concentration (left Y axis) and removal efficiency (right Y axis) of wastewater nutrients and organic carbon during the *Tetrasphaera*-enriched microbiome establishment process. (A) Orthophosphate (PO_4_^3−^-P), (B) Total organic carbon (TOC), (C) Ammonia nitrogen (NH_4_^+^-N), and (D) Total nitrogen (TN).

In stage I, the noticeable increase in NH_4_^+^-N removal and general decrease in P removal (between day 23 and day 46, see Fig. 1A against Fig. 1C) suggested an overall negative relationship between nitrification and the P removal process. Considering the high TN concentration in the effluent (Fig. 1D), the nitrification products (i.e., NO_2_^−^ and NO_3_^−^) may be accumulated, which could trigger the competition of denitrification for the organic carbon source. Therefore, the nitrification process should be controlled to promote PAO enrichment and enhance P removal. High P removal (85%) was achieved in only 17 days upon adding 1 mg/L allylthiourea in stage II, indicating that the growth and activity of PAOs were enhanced with the regulation of nitrification inhibitor. Although the possibility could not be excluded that the increase in N removal in the late phase of stage II could be partially attributed to the increase in the TOC/TN ratio (from 1.31 to 2.84), in stage IV, the TOC/TN ratio (2.90 ± 0.3) was stable, and both TN and NH_4_^+^-N removal increased after the transient shock and were maintained in the late phase. These results imply that the nitrification process of the microbial community recovered, contributing to the increase in N removal. After the long-term selection pressure of low dosage allylthiourea, the composition of the nitrifying bacteria in the EBPR microbiome could be shifted, which agreed with a previous study in that allylthiourea had an obvious effect on the structure of the nitrifying bacterial community [30].

### 3.2. The effect of allylthiourea on EBPR performance

To further investigate the effect of allylthiourea on the EBPR microbiome, batch experiments were conducted using 143-day microbiome biomass under different allylthiourea dosages (0 mg/L, 1 mg/L and 5 mg/L). Based on 16S rRNA gene amplicon sequencing analysis, the relative abundance of *Tetrasphaera* PAOs in the sampled 143-day EBPR microbiome was 30.3%. In addition, the corresponding removal efficiencies of PO_4_^3−^-P, TOC, NH_4_^+^-N, and TN in the mother EBPR reactor at 143 days were 91.9%, 97.9%, 92.3%, and 64.4%, respectively (Fig. 1). The P uptake rate with 1 mg/L allylthiourea addition was highest (0.07 mg/L·min), followed by 0 and 5 mg/L (0.04 and 0.05 mg/L·min, respectively) (Fig. 2A). In addition, the P removal efficiency in the 5 mg/L group was 67.9%, which was the lowest among the three groups (Fig. 2C). The NH_4_^+^-N profiles of the three groups were different, especially in the aerobic phase. NH_4_^+^-N removal did not occur under 5 mg/L allylthiourea, while a remarkable decline in NH_4_^+^-N concentration was observed with 0 mg/L and 1 mg/L allylthiourea. For example, the NH_4_^+^-N concentration decreased from 24.64 mg/L to 1.48 mg/L when 1 mg/L allylthiourea was added (Fig. 2B). In addition, the dynamics of dissolved oxygen (DO) showed no visible difference in the DO concentration among the three experimental groups in the anaerobic phase (< 0.2 mg/L), while in the following aerobic phase, the DO concentration increased to 5.21 mg/L at 53 mins when exposed to 5 mg/L allylthiourea (Fig. 2D). In comparison, the DO concentrations were 1.09 mg/L and 1.04 mg/L with allylthiourea concentrations of 0 mg/L and 1 mg/L, respectively.

**Fig. 2.**
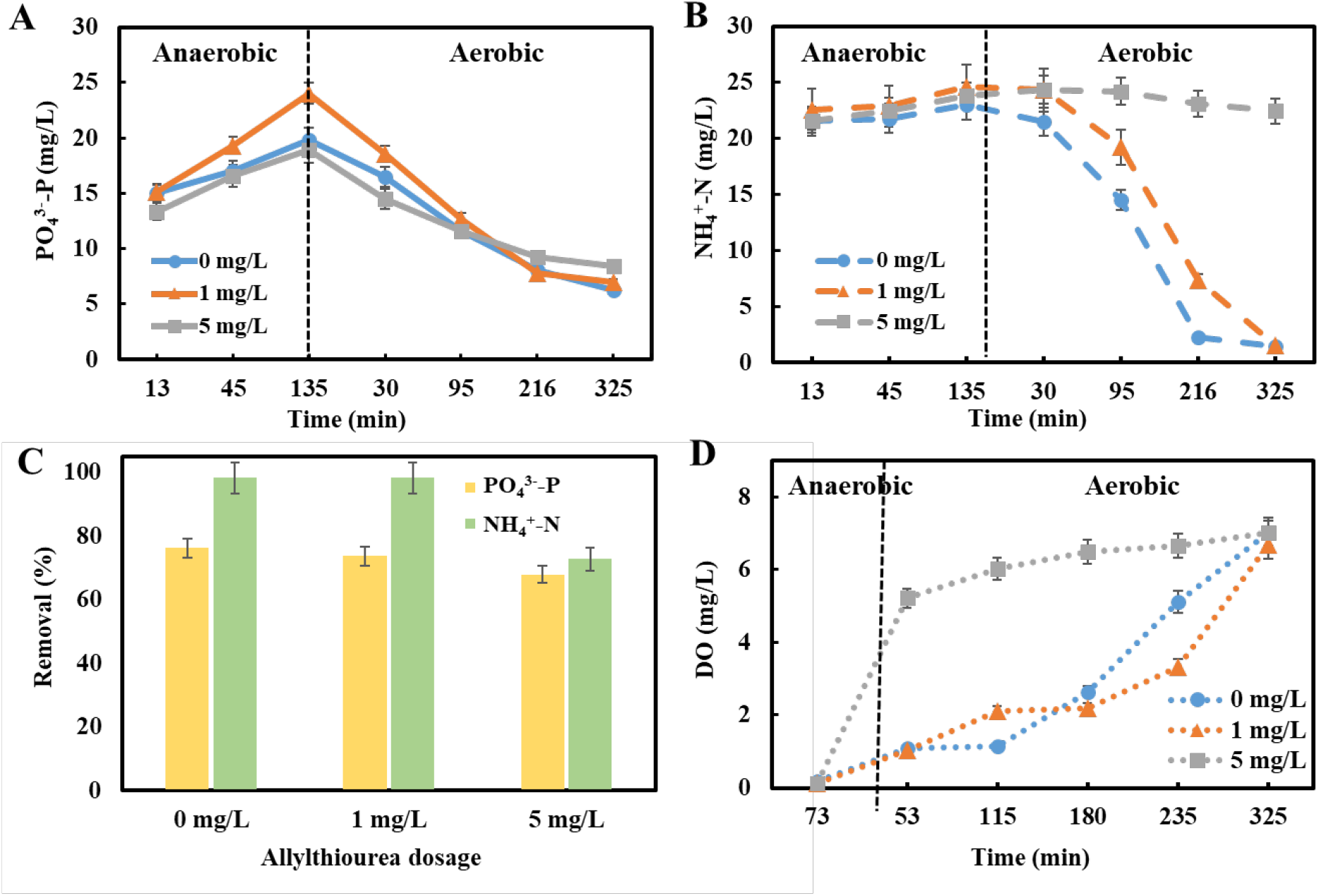
The profiles of nutrient dynamics and removal efficiency of the *Tetrasphaera*-enriched microbiome with different dosages of allylthiourea. Batch experiments were set up to comparatively test the effects of 0, 1, and 5 mg/L of allylthiourea. (A) PO_4_^3−^-P dynamics; (B) NH_4_^+^-N dynamics. (C) Removal efficiency of PO_4_^3−^-P and NH_4_^+^-N. (D) DO dynamics.

With the dosages of 0 and 1 mg/L allylthiourea, the enriched microbial community achieved stable N and P removal. In contrast, the high allylthiourea dosage (5 mg/L) showed a negative effect on the *Tetrasphaera*-enriched microbiome, consistent with the impact of allylthiourea addition on the nutrient removal performance previously observed in the EBPR reactor at stage III. The increased allylthiourea dosage (5 mg/L) inhibited the nitrification process, which could cause ammonia accumulation and disturbance in the microbial community [31]. In addition, when nitrification was completely inhibited, the production of NO_3_^−^ and NO_2_^−^ decreased, which may affect the activity and function of denitrifying PAOs. A previous study reported that although allylthiourea is soluble in water, it can be adsorbed by soil [30], indicating that allylthiourea may also be distributed in the sludge in addition to the aqueous phase. In addition, allylthiourea can be degraded by the microorganism; for example, a previous study indicated that 20 mg/L allylthiourea was degraded to 0.5–1.0 mg/L by activated sludge under aerobic conditions in 4 days [32], implying its low-security risk when applied for short-term biostimulation of wastewater treatment systems.

### 3.3. FISH analysis of the *Tetrasphaera*-enriched microbiome and evaluation of P recovery potential

The *Tetrasphaera*-enriched microbiome on day 143 was visualized by FISH. The targeted *Ca.* Accumulibacter and *Tetrasphaera* PAOs were visually detected in both the activated sludge sample (A) collected from the same WWTP as the seeding sludge (on December 2^nd^, 2020) and the *Tetrasphaera*-enriched microbiome (B) in our EBPR reactor (Fig. 3). Overall, *Tetrasphaera* PAOs were morphologically dominant in the EBPR microbiome, indicating that they were successfully enriched by the top-down design approach employed. Furthermore, the P storage ability and recovery potential of the *Tetrasphaera*-enriched microbiome were evaluated by determining the P content in the sludge ash using SEM‒EDS. The P content (counted as the weight percentage) in *Tetrasphaera*-enriched sludge ash and secondary sink sludge ash was 32.5% and 1.4%, respectively. Therefore, the percentage of P in *Tetrasphaera-*enriched microbiome sludge ash dramatically increased by 23.2 times compared with the normal full-scale WWTP sludge, indicating that the P-storage ability of the *Tetrasphaera*-enriched microbiome was largely enhanced. A strong signal intensity of Na and Mg elements was detected in the *Tetrasphaera*-enriched sludge, while the Al and Si signal intensities were higher in WWTP sludge (Fig. 3C). These results corresponded with the surface morphology of these two kinds of sludge (Fig. 3A and Fig. 3B). The WWTP sludge ash showed a red color with a loose structure because the WWTP influent contained sand, and the WWTP used Al-based coagulants to aid nutrient removal from wastewater. In comparison, the *Tetrasphaera*-enriched sludge ash showed a white color and compact morphology, which resulted from the designed operation process and influent compositions (e.g., MgSO_4_ and CaCl_2_). Theoretically, *Tetrasphaera*-enriched sludge ash is attractive for further use in white P recovery and is promising to replace the depleting phosphate rock, in which Ca_3_(PO_4_)_2_ is the main component [33]. Therefore, three essential conditions to promote P recovery from wastewater are achieved: i) high P removal efficiency, ii) microbiome with enhanced P storage ability, and iii) P-rich ashes and compounds for further value-added P recovery. The establishment of the *Tetrasphaera*-enriched microbiome holds promise for exploiting *Tetrasphaera* microbiology and EBPR biotechnology for the targeted regulation and promotion of biological P recovery in full-scale WWTPs.

**Fig. 3.**
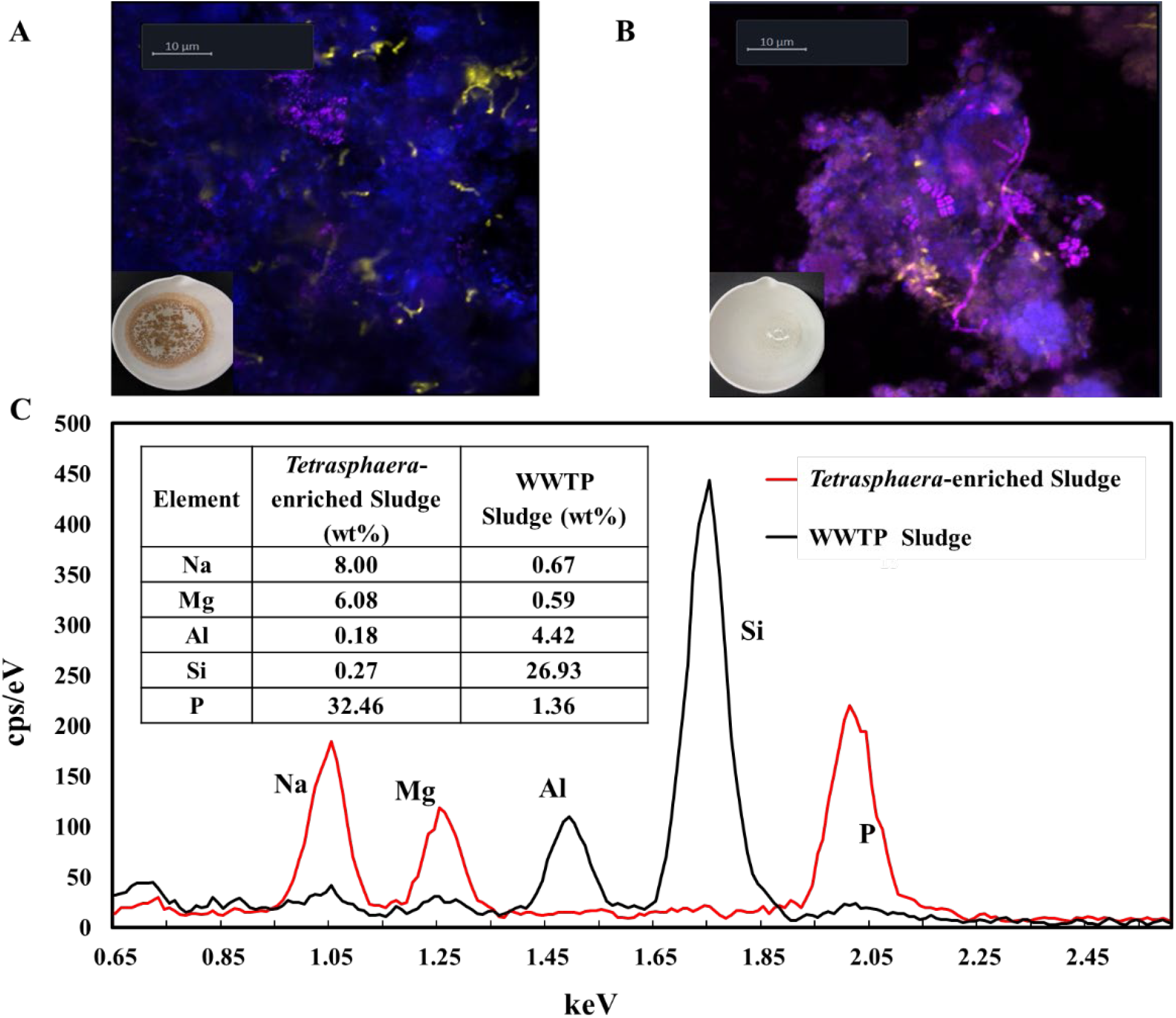
Fluorescence *in situ* hybridization (FISH) and elemental distribution analyses of full-scale WWTP sludge and lab-scale *Tetrasphaera*-enriched sludge. (A-B) FISH images of local WWTP sludge collected on December 2^nd^, 2020 (A) and the *Tetrasphaera*-enriched microbiome on day 143 (B). The colors denote stained bacterial fractions in the community, with all bacteria in blue, *Ca.* Accumulibacter is yellow, and *Tetrasphaera* is purple. (C) Elemental distribution and weight content (%) of *Tetrasphaera*-rich sludge ash and WWTP sludge ash by scanning electron microscopy and energy dispersive X-ray spectroscopy (SEM‒EDS) analysis.

### 3.4. The microbiome composition and dynamics during the establishment process of the *Tetrasphaera*-enriched microbiome

The microbiome composition and temporal dynamics during the enrichment process of *Tetrasphaera* PAOs were investigated based on high-throughput 16S rRNA gene amplicon sequencing (Fig. 4). In general, the microbiome was dominated by *Tetrasphaera* PAOs in coexistence with *Microlunatus* PAOs and glycogen accumulating organisms (GAOs) (i.e., *Ca.* Competibacter). In seed sludge, *Tetrasphaera* PAOs, *Microlunatus* PAOs, *Brevundimonas*, and *Ca.* Competibacter GAOs showed relative abundance of 1.5%, 0.1%, 0.3% and 12.4%, respectively. For the initial establishment of the EBPR microbiome, their relative abundance on day 73 shifted to 13.5%, 2.3%, 15.1%, and 3.0%, respectively. In addition, the highest relative abundance of *Tetrasphaera* PAOs was approximately 40% on day 113, approximately equivalent to a cell or biovolume percentage of 62% to 67% by assuming an average 16S rRNA gene copy number of 1.0 for *Tetrasphaera* and between 2.5 and 3.0 for activated sludge bacteria [34]. This result implied that allylthiourea promoted significant enrichment of *Tetrasphaera* PAOs (*P value* = 0.017, Kruskal-Wallis test, Fig. S1A). Notably, the *Tetrasphaera* PAOs were the most abundant 16S rRNA gene ASV in the EBPR microbiome (i.e., EBPR-ASV0001). Phylogenetic analysis showed that *Tetrasphaera sp.* EBPR-ASV0001 was phylogenetically different from the six previously reported isolates (Fig. 4B and Table S3). The global similarity of 16S rRNA gene sequences between EBPR-ASV0001 and *Tetrasphaera japonica* was 96.8%, indicating that EBPR-ASV0001 is most likely a denitrifying PAO similar to *T. japonica*.

**Fig. 4.**
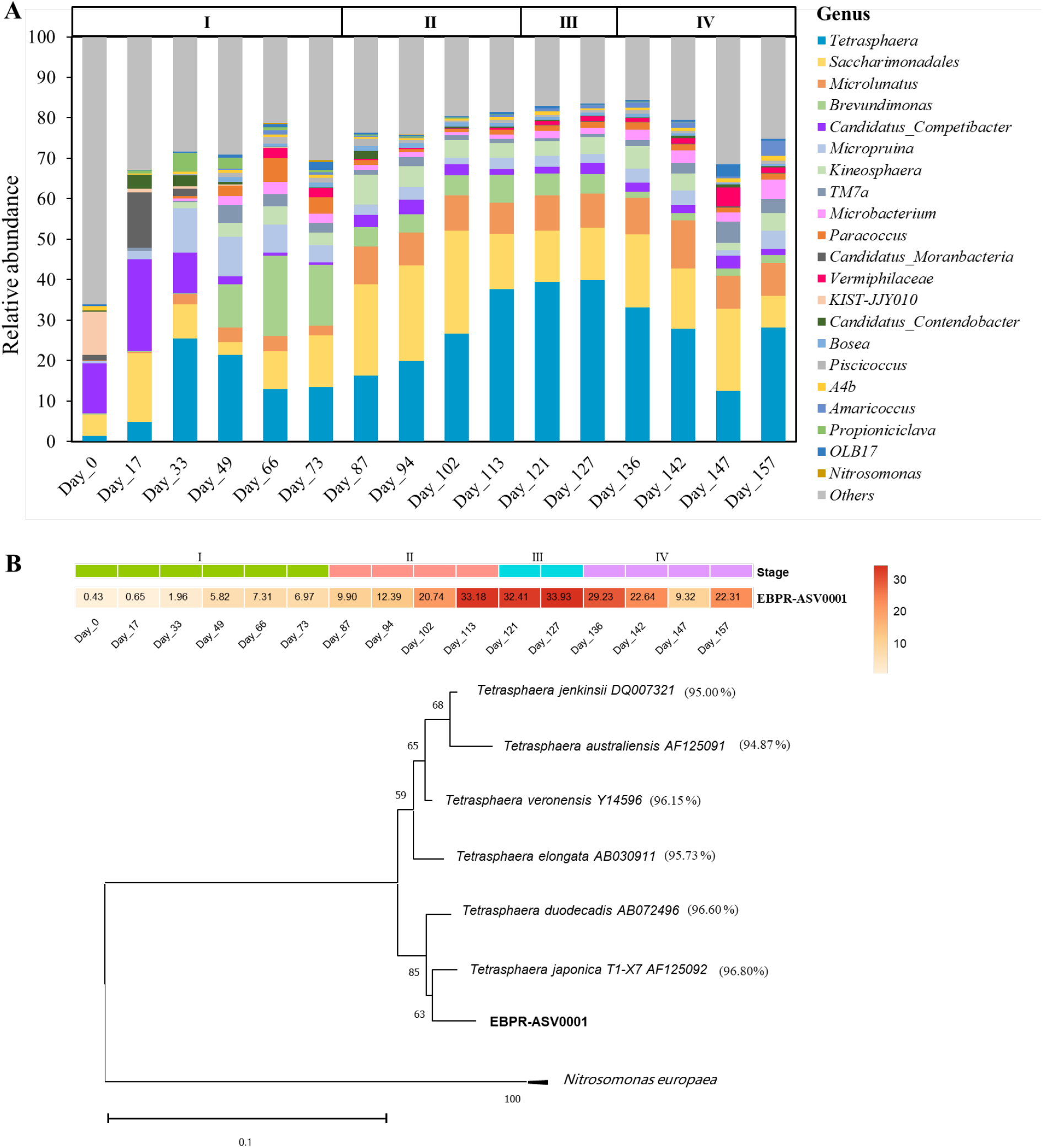
Microbial composition and dynamics during the establishment process of the *Tetrasphaera*-enriched microbiome. (A) The top 20 bacteria at the genus level; (B) The dynamic and phylogenetic analysis of the most abundant *Tetrasphaera*-related bacteria (EBPR-ASV0001). The evolutionary history was inferred by the maximum likelihood method and Kimura 2-parameter model. Two *Nitrosomonas europaea* sequences were used as the outgroup. The global similarity of EBRP-ASV0001 compared with six *Tetrasphaera* isolates is provided within the brackets.

The abundance fluctuations of *Ca.* Competibacter GAOs, *Brevundimonas*, and *Tetrasphaera* PAOs during stage I (day 0-73, Fig. 4A) were consistent with unstable biological phosphorus and nitrogen removal. Specifically, *Ca.* Competibacter GAOs decreased significantly in the first 33 days (*P value*=0.05, Kruskal-Wallis test), while *Brevundimonas* and *Paracoccus* were markedly enriched during days 49-73 (*P value*=0.05, Kruskal-Wallis test) (Fig. S1B, C, and D). In addition, the decrease in the relative abundance of *Tetrasphaera* PAOs and the P removal efficiency coincided with the prosperity of *Brevundimonas* and *Paracoccus*. Previous reports indicated that the occurrence of the denitrification process in the anaerobic phase could cause EBPR failure because some denitrifiers could compete for organics with PAOs [16]. *Brevundimonas* and *Paracoccus* are well known not only as denitrifiers but also as heterotrophic nitrification and aerobic denitrification (HN-AD) bacteria, which can simultaneously perform nitrification and denitrification under aerobic conditions [35–37]. Therefore, their abundance increase could lead to potential competition for organic molecules and/other resources with *Tetrasphaera* PAOs, especially in full-scale WWTPs where the dissolved oxygen is heterogeneous.

With the addition of 1 mg/L allylthiourea, the following gradual abundance increase in *Tetrasphaera* PAOs (13.0% to 37.7%) was coupled with the sharp abundance decrease in *Nitrosomonas* AOBs (0.35% to 0.02%) as well as the denitrifiers *Brevundimonas* (19.9% to 6.9%) and *Paracoccus* (5.8% to 1.1%) from day 66 (stage I) to day 113 (stage II). This result can be explained by the chain effects associated with the allylthiourea inhibition of nitrifying *Nitrosomonas*, i.e., the inhibition of autotrophic ammonia oxidation largely reduced denitrifying substrates available to the dentritrifers, thus greatly alleviating their competition against PAOs for organic resources (Fig. 4A). Interestingly, the ammonia oxidizer *Nitrosomonas* decreased by 13.7 times from days 73 to 113 and disappeared on day 121, while high ammonia removal was still observed (e.g., 98.7% on day 121, Fig. 1C), implying the existence of an alternative nitrification process. We speculated that some heterotrophic nitrifiers should participate in the nitrification process when autotrophic nitrifiers (e.g., *Nitrosomonas*) are evidently inhibited under a low dosage of allylthiourea (i.e., 1 mg/L), facilitating the nitrogen removal. In stage III, when the transient shock of 5 mg/L allylthiourea was applied, PAOs fluctuated in their relative abundance, which first decreased to 12.6% on day 147 and then increased to 28.2% on day 157. We hypothesized that a high allylthiourea dosage would adversely affect the proliferation and function of denitrifying PAOs that can couple nitrate and/or nitrite reduction to P uptake due to the heavy suppression of nitrification. Supporting this hypothesis, ammonia removal was completely suppressed in stage III, accompanying a dramatic decline in the removal efficiency of TN and PO_4_^3−^-P to 19.5% and 58.4%, respectively (Fig. 1A). Furthermore, a high allylthiourea dosage restrained the propagation of the most abundant PAOs, *Tetrasphaera* sp. EBPR-ASV0001, which showed a marked decline on day 142 (22.64%, Fig. 1B). Further genome-based study and laboratory isolation of *Tetrasphaera* sp. EBPR-ASV0001 to reveal its metabolic pathways, physiology, and application potential in full-scale EBPR WWTPs are warranted.

### 3.5. The microbial co-occurrence patterns during the establishment process of the *Tetrasphaera-*enriched microbiome

To predict microbial interactions in the *Tetrasphaera*-enriched microbiome, microbial co-occurrence patterns over the 170-day reactor operation were analyzed based on network analysis (Fig. 5). The positive bacterial association network had 85 nodes (ASVs) and 167 edges (correlations, Table S4), which were topologically partitioned into nine modules presumably representing discrete and interactive ecological niches over time (Fig. 5A). The topological properties of the observed network were calculated and compared with those of identically sized Erdös-Réyni random networks (Table S4). The clustering coefficient (CC) and modularity index of the observed *Tetrasphaera*-enriched co-occurrence network were 0.449 and 0.687 and higher than those of the corresponding Erdös-Réyni random networks (0.046 and 0.418), suggesting that the microbiome assembly process is nonrandom (or deterministic). Moreover, the network of the *Tetrasphaera*-enriched microbiome showed small-world properties with a coefficient 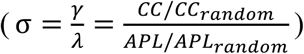 [38] of 8.97, suggesting that the enrichment strategy adjusts the overall microbial interconnections and interior interaction relationship.

**Fig. 5.**
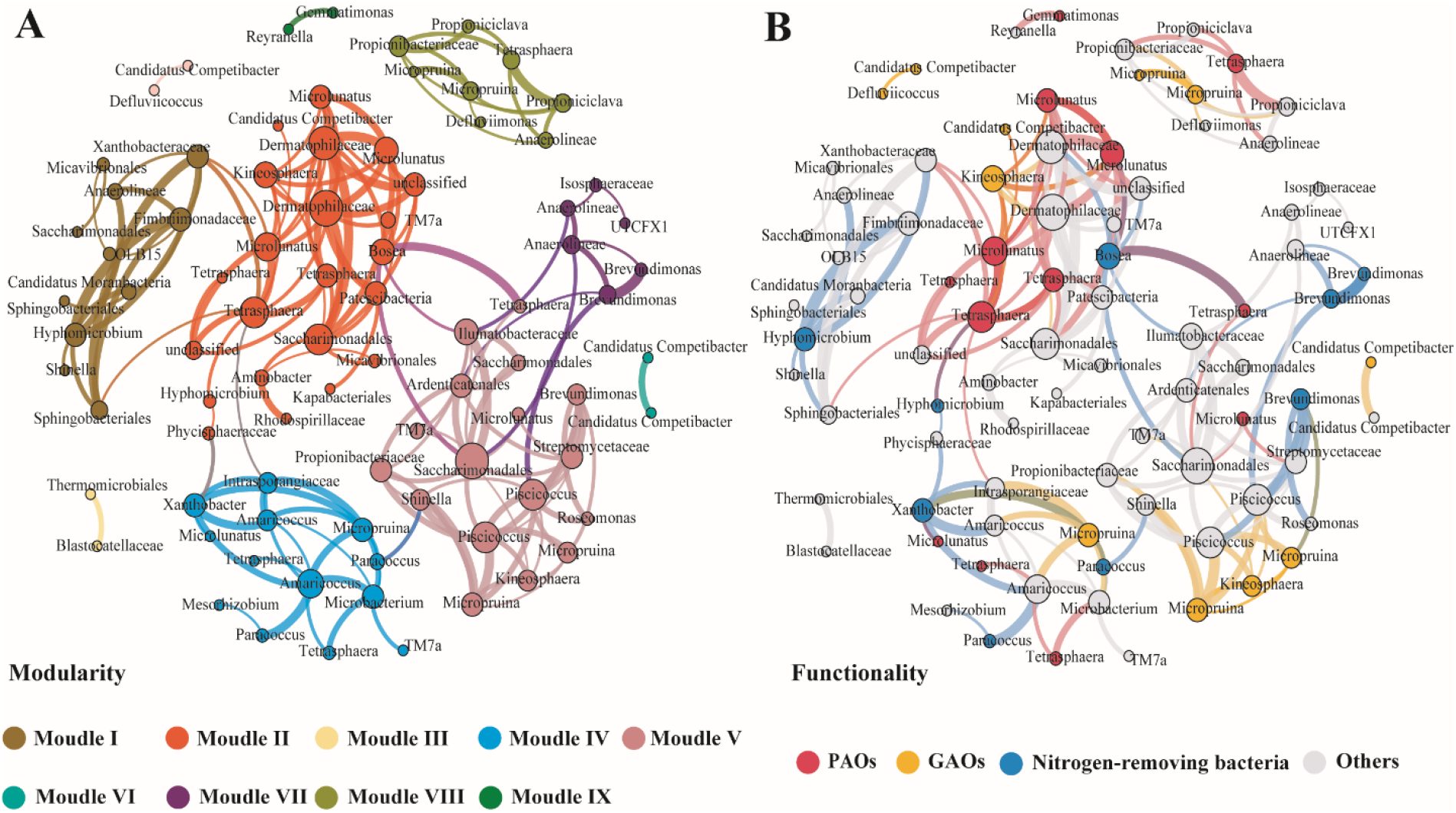
Co-occurrence network of the *Tetrasphaera*-enriched microbiome-wide associations based on pairwise Spearman’s correlations between bacterial ASVs. Each connection (i.e., edge) stands for a significant Spearman’s correlation coefficient > 0.6 and a *P value* < 0.01. The size of each node is proportional to the number of connections (i.e., degree), and the thickness of the edge is scaled according to the strengths of the correlation between nodes. (A) Co-occurrence network colored by modularity class. (B) Co-occurrence network colored by functionality. PAOs: *Tetrasphaera*, *Microlunatus*, and *Gemmatimonas*; GAOs: *Micropruina*, *Kineosphaera*, and *Candidatus* Competibacter; Nitrogen-removing bacteria: *Xanthobacter*, *Paracoccus*, *Hyphomicrobium*, *Brevundimonas*, and *Bosea*.

To link ecological niches of co-occurring taxa to their functionality, each network node was taxonomically assigned to three functional groups (Fig. 5B), according to previous reports [7, 29, 36, 39], including PAOs, GAOs, and nitrogen-removing bacteria (e.g., denitrifiers and HN-AD bacteria). Overall, *Tetrasphaera* was distributed in four modules of the network (i.e., module II, IV, V, and VIII), implying niche differences between members of the genus *Tetrasphaera*. This observation supported the previous conclusion [4, 10, 12–14], namely, the different species of *Tetrasphaera* could have a specific metabolic preference; therefore, the genus *Tetrasphaera* was extremely versatile and could widely interact with coexisting bacteria in EBPR. In particular, *Tetrasphaera* PAOs significantly co-occurred with *Microlunatus* PAOs in module II (Spearman’s ρ =0.906, FDR-adjusted *P* < 0.01, Fig. 5), indicating their potential cooperative relationships. In addition, known denitrifiers (e.g., *Hyphomicrobium, Bosea*, and *Xanthobacter*) occurred with GAOs (e.g., *Ca.* Competibacter, *Kineosphaera*, and *Micropruina*) in modules I, II and IV, implying a shared niche preference, such as taking up organic matter for growth. The wide within-module co-occurrence between denitrifiers and organic-degrading bacteria (e.g., *Hyphomicrobium* and *Shinella* in module I, *Bosea* and *TM7a* in module II, and *Xanthobacter* and *Amaricoccus* in module IV) supports the classical presumption of their cooperative interactions, such as cross-feeding, i.e., the former provides small organic molecules for the latter as a carbon source.

The microbial interaction and niche patterns revealed in the co-occurrence network of the *Tetrasphaera*-enriched microbiome (Fig. 5) were associated with the dynamics of the removal efficiency of nutrients (A, C, and D) rather than organic carbon (B, Fig. 1). First, the wide co-occurrence and prevalence of denitrifiers and organic-degrading bacteria in the same modules, together with the near-perfect TOC removal (95±1.2%) and mostly favorable TN removal (77±2.9% when no inhibition occurred), implies that chemoorganotrophic respiration and heterotrophic denitrification could be active during the microbiome assembly process. Moreover, *Micropruina* was a fermentative GAO [4], and its positive correlations with *Brevundimonas* in module V (Spearman’s ρ = 0.816) and *Paracoccus* in module IV (Spearman’s ρ = 0.818) indicated the fermentative bacteria may cross-feed co-occurring HN-AD bacteria with fermentation products. Likewise, a positive correlation between *Tetrasphaera* PAOs and *Bosea* denitrifiers (Spearman’s ρ = 0.910, Fig. 5) suggests that denitrifying phosphorus removal should have occurred in the reactor, promoting simultaneous biological N and P removal. A prior genomic survey indicated that *Tetrasphaera* can utilize nitrite for poly-P production [11], which could explain the abovementioned enrichment scenarios of *Tetrasphaera* PAOs facilitated by cross-feeding partnerships with nitrate denitrifiers. Therefore, the enrichment strategy with intricately designed multicarbon sources and low-dose allylthiourea creates microbial niches and fosters diverse cooperative interactions that affect the microbiome composition and establishment.

### 3.6. The nexus between microbiome, environment, and EBPR performance

In addition to microbiome dynamics and co-occurrence patterns, the roles of abiotic factors (i.e., environmental condition and influent composition) and biotic factors (i.e., microbiome composition and interaction) in driving *Tetrasphaera*-enriched microbiome establishment were explored based on RDA and PCoA analyses. The microbial communities shifted away upon reactor startup and quickly assembled with the continuous feedings of synthetic wastewater containing multicarbon sources (S1, Fig. 6, and Fig. S2). When the nitrification inhibitor (i.e., allylthiourea) was introduced, the microbial community was significantly selected and restructured (S2, S3, and S4, Fig. 6 and Table S5). The nexus between the environmental condition and microbial community was visualized with the first two RDA axes (Fig. 6A), which jointly explained 52.2% of the total microbiome variances and indicated that the designed environmental parameters, especially the inhibitor (adjusted R^2^ = 0.23, *P value* < 0.01, Table S6), were among the major drivers of the microbiome assembly process. In addition, the *Tetrasphaera* PAOs and *Microlunatus* PAOs positively correlated with allylthiourea and PO_4_^3−^-P, indicating that *Tetrasphaera* PAOs and *Microlunatus* PAOs have growth advantages under selective conditions. The association analysis between the microbial community and EBPR performance showed that *Tetrasphaera* PAOs, *Microlunatus* PAOs, and *Saccharimonadales* positively correlated with PO_4_^3−^-P, TN, and TOC removal. Meanwhile, *Brevundimonas* and *Kineosphaera* were positively correlated with NH_4_^+^-N removal (Fig. 6B). Consistent with the RDA results, Mantel test analysis revealed that allylthiourea significantly affected the microbiome dynamics and PO_4_^3−^-P removal (*P value* < 0.005, Table S7).

**Fig. 6.**
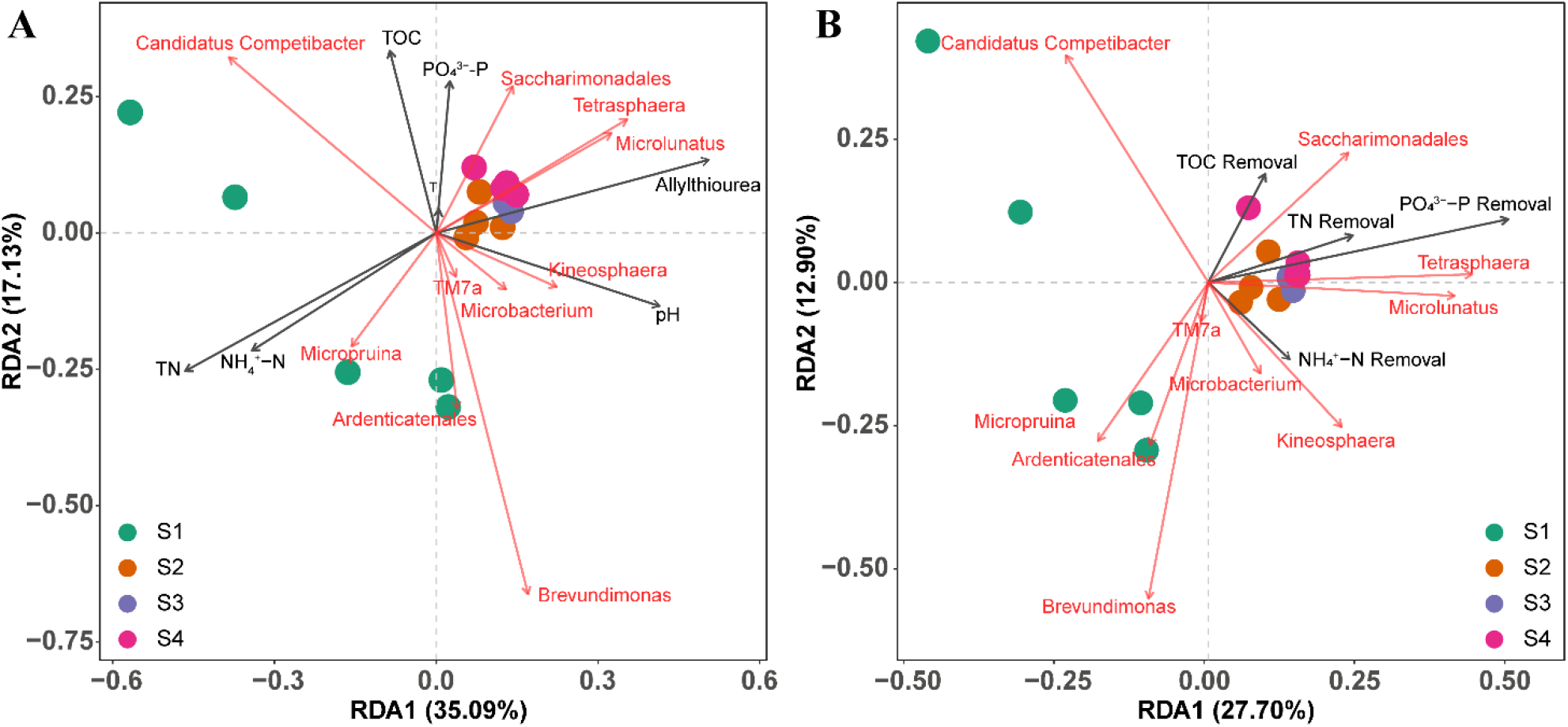
Redundancy analysis (RDA) of (A) the nexus between the microbial community and environmental variables and (B) the nexus between the microbial community and EBPR performance. Four operational stages were defined based on the usage of nitrification inhibitor allylthiourea during the microbiome establishment process: S1: Natural enrichment phase (day 0-73) with no inhibitor; S2: Reinforced enrichment phase (day 74-120) with 1 mg/L allylthiourea as inhibitor; S3: Transient shock phase (day 121-127) with 5 mg/L allylthiourea as inhibitor); S4: Recovery phase (day 128-170) with allylthiourea dosage adjusted back to 1 mg/L. The dashed arrow lines denote two adjacent sampling time points in the first 73 days after reactor startup.

Overall, the correlation nexus between microbial (M) community, environmental (E) variables, and EBPR performance (P) (defined here as the ‘MEP nexus’) indicated that 1 mg/L allylthiourea, as a key abiotic factor, promoted the selection and proliferation of *Tetrasphaera* PAOs, *Microlunatus* PAOs and organic degrading bacteria *Saccharimonadales* while inhibiting *Nitrosomonas* AOBs and indirectly affecting potential competitors such as denitrifying *Brevundimonas*. Furthermore, the removal of carbon and nutrients (i.e., TOC, TN, and PO_4_^3−^-P) was enhanced. Notably, *Tetrasphaera* and *Microlunatus* PAOs showed no significant negative correlation in their relative abundance in the whole process (Fig. S3), and their positive co-occurrence patterns (Module II, Fig. 5A) made their coexistence feasible in the *Tetrasphaera*-enriched EBPR microbiome. In addition, *Saccharimonadales* is reported to be a key degrader of organic molecules (e.g., glucose and sugars) in aerobic sludge systems [40]. Based on their temporal dynamics, *Microlunatus* PAOs and *Saccharimonadales* should be the key biotic factors that benefit from the designed multicarbon sources and positively contribute to building a *Tetrasphaera*-enriched microbiome and enhancing the reactor performance for organics and P removal. In summary, the MEP nexus indicates that the enrichment strategy designed here has realized the selective enrichment of *Tetrasphaera* PAOs as an outcome of co-actions from both abiotic and biotic factors.

## 4. Conclusion

In this study, an enrichment strategy featuring intricately designed multicarbon sources amended with low dosage allylthiourea was proposed and applied to build a *Tetrasphaera*-enriched EBPR microbiome. Based on 16S rRNA gene amplicon sequencing analysis, a novel putative *Tetrasphaera* PAO species, EBPR-ASV0001, was identified and enriched at a relative abundance of 40%. The established microbiome showed enhanced P removal (~85%) and N removal (~80%) and a 23.2 times higher P content in the sludge ash than in the normal full-scale WWTP sludge. The addition of low-dosage allylthiourea (e.g., 1 mg/L demonstrated in this study) regulated the community structure and affected the microbial co-occurrence patterns, which can be advised as a short-term biostimulation approach for the quick selection and enrichment of *Tetrasphaera* PAOs to strengthen biological P removal. The findings of this study expand our knowledge of the temporal dynamics and interaction patterns of a novel putative PAO member of *Tetrasphaera*. Future studies with metagenomic analysis or full-length 16S rRNA sequencing of the enriched *Tetrasphaera* species are needed to verify their phylogeny and putative function more reliably as new PAOs. Nonetheless, from an engineering perspective, this outcome provides theoretical guidance for exploiting and optimizing EBPR application for sustainable P removal and recovery coupled with biological nitrogen removal from high-concentration wastewater.

## Acknowledgement

This work was supported by the Key R&D Program of Zhejiang (2022C03075), Zhejiang Provincial Natural Science Foundation of China under Grant No. LR22D010001. We would like to thank Dr. Xiao Yang and Dr. Xiangyu Yang for the helpful discussion and technical advice. The author would like to thank Yisong Xu for her professional support in equipment procurement and lab management. We thank the Microscopy Core Facility of Westlake University for the facility support and thank technician Fang Xiao for technical assistance. We thank the Research Center for Industries of the Future (RCIF), the Instrumentation and Service Center for Molecular Sciences and Physical Sciences, and The Westlake University-Muyuan Group Joint Research Institute at Westlake University for support. We thank the Westlake University HPC Center for computation support.

## Data availability

The raw data of 16S rRNA gene amplicon sequencing were deposited in China National GeneBank (CNGB) under the project accession number CNP0002991.

