## Supplementary for "Temporal Dynamics and Performance Association of the *Tetrasphaera*-Enriched Microbiome for Enhanced Biological Phosphorus Removal"

**Legend of SI:**

**Text S1** Bioreactor operation and monitoring

**Text S2** Fluorescence *in situ* hybridization (FISH) experiment

**Text S3** Co-occurrence network construction

**Table S1.** The information of stock solutions used to prepare the synthetic wastewater in this study.

**Table S2.** The FISH probe used for the detection of different PAO groups in this study.

**Table S3.** The BLAST results showing the top 20 matches of EBRP-ASV0001* in the

NCBI's nt (nucleotide collection) database. The V1-V3 regions of 16S rRNA gene sequence of EBRP-ASV0001 showed a global similarity between 95.7% and 96.8% to six isolates of *Tetrasphaera*.

**Table S4.** Topological properties of the observed co-occurrence network of the *Tetrasphaera*-enriched microbiome and the corresponding identically-sized Erdös-Réyni random network.

**Table S5**. Permutation Based Analysis of Variance (PERMANOVA) test result based on PCoA results. S1: Natural enrichment phase (day 0-73) with no inhibitor; S2: Reinforced enrichment phase (day 74-120) with 1 mg/L allylthiourea as inhibitor; S3: Transient shock phase (day 121-127) with 5 mg/L allylthiourea as inhibitor); S4: Recovery phase (day 128-170) with allylthiourea dosage adjusted back to 1 mg/L.

**Table S6.** The relationship between environment condition and microbial community. The significance of each environment condition effect was tested with the ANOVA based on the RDA results.

**Table S7.** Mantel tests of correlations between microbial communities, environmental variables, and EBPR performance. EBPR: enhanced biological phosphorus removal.

**Fig. S1** Statistics test of (A) dynamic of *Tetrasphaera* before (day 0-73) and after (day 87-157) allylthiourea addition (B) dynamic of *Candidatus* Competibacter in day 0-33 and day 49-73. (C) dynamic of *Brevundimonas* in day 0-33 and day 49-73. (C) dynamic of *Paracoccus* in day 0-33 and day 49-73.

**Fig. S2** Principal coordinate analysis (PCoA) plot showing the *Tetrasphaera*-enriched microbiome establishment process. Four operational stages were defined based on the usage of nitrification inhibitor allylthiourea during the microbiome establishment process: S1: Natural enrichment phase (day 0-73) with no inhibitor; S2: Reinforced enrichment phase (day 74-120) with 1 mg/L allylthiourea as inhibitor; S3: Transient shock phase (day 121-127) with 5 mg/L allylthiourea as inhibitor); S4: Recovery phase (day 128-170) with allylthiourea dosage adjusted back to 1 mg/L.

**Fig. S3** The microbial dynamic of *Tetrasphaera* PAOs and *Microlunatus* PAOs during the whole establishment process of the *Tetrasphaera*-enriched microbiome.

**Text S1** Bioreactor operation and monitoring

In each cycle, 5-litre of synthetic wastewater (see the formula in Table S1) was fed to the reactor, and the effluent was regularly collected for chemical analyses. The reactor was operated at a room temperature between 20 °C and 24 °C, a mechanic mixing speed of 150 rpm, a hydraulic retention time (HRT) of 16 h, and a sludge retention time (SRT) of 12 d. The total solids (TS) and volatile solids (VS) content in the reactor were maintained at 6.0 ± 1.0 g/L and 5.5 ± 1.0 g/L, respectively. The dissolved oxygen (DO) concentration, oxidation-reduction potential (ORP), and pH were continuously monitored online using an LDO II sensor (Hach), an ORP probe (GLI3/4, Hach), and a pH probe (GLI3/4, Hach), respectively. Nitrogen gas was spared through the reactor at a flow rate of about 1 L/min during the anaerobic period. The pH was controlled at 7.2 ± 0.2 by the addition of 1 M hydrochloric acid and 1 M sodium hydroxide stock solutions. The air was supplied at a flow rate of about 2 L/min to maintain DO concentration reached 2 ± 0.2 mg/L during aerobic period.

**Text S2** Fluorescence *in situ* hybridization (FISH) experiment

Briefly, the biomass collected from the reactor were fixed by adding 37% formaldehyde solution (the ratio between biomass and formaldehyde is 10:1) and then held at 4 °C for 2 hours. After centrifugation followed by a twice wash (suspending samples in the mix solution of phosphate buffered saline (PBS) and ethanol (volume 1:1)), the resuspended biomass was collected and then loaded into the glass slides for incubation. The incubation condition was 65 °C for 1 hour, and then the slides were immersed in 50% ethanol, 80% ethanol, and 100% ethanol in turn for dehydration. After drying in weak airflow, the probes were loaded with prepared hybridization buffer (35% formamide) for hybridization. The hybridization condition was 46 °C for 12 hours. The EUBmix probe that composed of three probes (EUB338-1, EUB338-2, and EUB338-3) was used to target the entire bacterial community. To characterize the PAOs community characterization, the mixed *Ca.* Accumulibacter PAOs probe (Acc-I-444, Acc-II-444, and PAO651) and the mixed *Tetrasphaera* PAOs probe (Tet1-266, Tet2-892, Tet2-174, and Tet3-654) were used. The detailed information about the FISH probes can be found in Table S2. After washing using the prepared wash buffer followed by air drying, the probe labeled biomass was examined using a laser scanning confocal microscope (Zeiss, Zeiss LSM800, Germany).

**Text S3** Co-occurrence network construction

The ASVs matrix table was used to calculate the strong pairwise correlations between each node (Spearman's $\rho > 0.6$ and FDR-adjusted $P < 0.01$). The ASVs that occurred in at least eight samples were used for network analysis. Moreover, the topological features including modularity, clustering coefficient, average path length, average degree, and network diameter of the network were calculated. Meanwhile, 10 000 Erdös-Réyni random networks with the same number of nodes and edges as the real networks were generated, and a set of topological properties was calculated using the 'Random_network.R' script of MbioAssy1.0 (https://github.com/emblab-westlake/MbioAssy1.0) [1,2].

**Table S1.** The information of stock solutions used to prepare the synthetic wastewater in this study.

| Stock solution | Stock solution composition | Mass concentration in stock solution(g/L) | Taking volume (L) for the synthetic wastewater preparation (5L) |
|---|---|---|---|
| Solution A | Amicase | 35.29 | 0.06 (±0.03) |
| | Glucose | 9.37 | 0.15 (±0.1) |
| | Sodium acetate | 29.08 | 0.08 (±0.005) |
| Solution B | $K_2HPO_4$ | 5.96 | 0.06 (±0.005) |
| | $KH_2PO_4$ | 4.674 | 0.06 (±0.005) |
| Solution C | $NH_4Cl$ | 7.4 | 0.29 (±0.005) |
| | $MgSO_4 \cdot 7H_2O$ | 11.9 | 0.29 (±0.005) |
| | $CaCl_2 \cdot 2H_2O$ | 5.5 | 0.29 (±0.005) |

**Table S2.** The FISH probe used for the detection of different PAO groups in this study.

| Primer | Sequence (5' to 3') | Target group | Label | Ref. |
|---|---|---|---|---|
| EUB338-1 | GCTGCCTCCCGTAGGAGT | Many but not all Bacteria | 5`6-FAM | 1 |
| EUB338-2 | GCAGCCACCCGTAGGTGT | Planctomycetales | 5`6-FAM | 1 |
| EUB338-3 | GCTGCCACCCGTAGGTGT | Verrucomicrobiales | 5`6-FAM | 1 |
| Acc-I-444 | CCCAAGCAATTTCTTCCCC | *Candidatus* Accumulibacter Clade IA and other Type I clades | 5`Cy3 | 1 |
| Acc-II-444 | CCCGTGCAATTTCTTCCCC | *Candidatus* Accumulibacter Clade IIA, IIC and IID | 5`Cy3 | 1 |
| PAO651 | CCCTCTGCCAAACTCCAG | *Candidatus* Accumulibacter | 5`Cy3 | 1 |
| Tet1-266 | CCCGTCGTCGCCTGTAGC | Clone ASM31 | 5`Cy5 | 2 |
| Tet2-892 | TAGTTAGCCTTGCGGCCG | Clone ASM47 | 5`Cy5 | 2 |
| Tet2-174 | GCTCCGTCTCGTATCCGG | *T. jenkinsii*, *T. australiensis*, *T. veronensis*, and *Candidatus* N. limicola | 5`Cy5 | 2 |
| Tet3-654 | GGTCTCCCCTACCATACT | Unclutured *Tetrasphaera* | 5`Cy5 | 2 |

**Table S3.** The BLAST results showing the top 20 matches of EBRP-ASV0001* in the NCBI's nt (nucleotide collection) database. The V1-V3 regions of 16S rRNA gene sequence of EBRP-ASV0001 showed a global similarity between 95.7% and 96.8% to six isolates of *Tetrasphaera*.

| Sequence Name | Accession Number in NCBI | Similarity (%) |
|---|---|---|
| uncultured bacterium partial 16S rRNA gene | LR638470.1 | 98.08 |
| uncultured bacterium partial 16S rRNA gene | LR634716.1 | 98.08 |
| uncultured bacterium partial 16S rRNA gene | LR653116.1 | 98.08 |
| uncultured bacterium partial 16S rRNA gene | LR646250.1 | 98.08 |
| uncultured bacterium partial 16S rRNA gene | LR638410.1 | 97.86 |
| uncultured bacterium partial 16S rRNA gene | LR646970.1 | 97.86 |
| Uncultured Actinobacteria bacterium 16S rRNA gene from clone QEEB3BE04 | CU918095.1 | 97.86 |
| Uncultured Actinobacteria bacterium 16S rRNA gene from clone QEEB3AC11 | CU917872.1 | 97.86 |
| uncultured bacterium partial 16S rRNA gene | LR638621.1 | 97.65 |
| uncultured bacterium partial 16S rRNA gene | LR636792.1 | 97.44 |
| Uncultured bacterium clone F_SBR_6 16S ribosomal RNA gene, partial sequence | HQ010781.1 | 97.44 |
| uncultured bacterium partial 16S rRNA gene | LR635747.1 | 97.44 |
| uncultured bacterium partial 16S rRNA gene | LR639645.1 | 97.22 |
| uncultured bacterium partial 16S rRNA gene | LR636158.1 | 97.22 |
| uncultured bacterium partial 16S rRNA gene | LR634453.1 | 97.22 |
| uncultured bacterium partial 16S rRNA gene | LR654786.1 | 97.22 |
| uncultured bacterium partial 16S rRNA gene | LR653251.1 | 97.22 |
| uncultured bacterium partial 16S rRNA gene | LR651403.1 | 97.22 |
| uncultured bacterium partial 16S rRNA gene | LR650603.1 | 97.22 |
| Uncultured bacterium clone ncd1007g03c1 16S ribosomal RNA gene, partial sequence | HM341171.1 | 97.22 |
| Uncultured bacterium 16S rRNA gene, clone cD4807 | AJ617850.1 | 97.22 |
| ***Tetrasphaera japonica strain T1-X7*** | **AF125092** | **96.80** |
| ***Tetrasphaera australiensis strain 109*** | **AF125091** | **94.87** |
| ***Tetrasphaera jenkinsii strain Ben 74*** | **DQ007321** | **95.00** |
| ***Tetrasphaera veronensis*** | **Y14596** | **96.15** |
| ***Tetrasphaera duodecadis*** | **AB072496** | **96.60** |
| ***Tetrasphaera elongata*** | **AB030911** | **95.73** |

*The sequence of EBRP-ASV0001 is

GACGAACGCTGGCGGCGTGCTTAACACATGCAAGTCGAACGGTGACGAGGGAGCTTGCTCCTTCTGA
TCAGTGGCGAACGGGTGAGTAACACGTGAGTAACCTGCCCCAGACTCTGGAATAACCTCGGGAAACC
GGGGCTAATACCGGATACGAGACGAAGAGGCATCTCTATCGTCTGGAAAGTTTTTCGGTCTGGGATGG
ACTCGCGGCCTATCAGCTTGTTGGTGAGGTAACGGCTCACCAAGGCGACGACGGGTAGCCGGCCTGA
GAGGGCGATCGGCCACACTGGGACTGAGACACGGCCCAGACTCCTACGGGAGGCAGCAGTGGGGAA
TATTGCACAATGGGCGAAAGCCTGATGCAGCGACGCCGCGTGAGGGATGACGGCCTTCGGGTTGTAA
ACCTCTTTCAGCAGGGAAGAAGCGCAAGTGACGGTACCTGCAAAAGAAGCACCGGCTAACTACGTG

**Table S4.** Topological properties of the observed co-occurrence network of the *Tetrasphaera*-enriched microbiome and the corresponding identically-sized Erdös-Réyni random network.

| | *Tetrasphaera*-enriched microbiome network | The corresponding random network [a] |
|---|---|---|
| Nodes | 85 | 85 |
| Edges | 167 | 167 |
| Modularity (MD) | 0.687 | 0.418 ±0.026 |
| Clustering Coefficient (CC) | 0.449 | 0.046 ± 0.013 |
| Average path length (APL) | 3.609 | 3.315 ±0.062 |
| Average degree (AD) | 3.929 | 3.929 |
| Network diameter (ND) | 11 | 7 |
| Graph density (GD) | 0.047 | 0.047 |

**Table S5**. Permutation Based Analysis of Variance (PERMANOVA) test result based on PCoA results. S1: Natural enrichment phase (day 0-73) with no inhibitor; S2: Reinforced enrichment phase (day 74-120) with 1 mg/L allylthiourea as inhibitor; S3: Transient shock phase (day 121-127) with 5 mg/L allylthiourea as inhibitor); S4: Recovery phase (day 128-170) with allylthiourea dosage adjusted back to 1 mg/L.

| Groups | $R^2$ | p-value | Significance |
|---|---|---|---|
| **S1 vs S2** | **0.33** | **0.011** | * |
| **S1 vs S3** | **0.29** | **0.007** | ** |
| **S1vs S4** | **0.35** | **0.009** | ** |
| S2 vs S3 | 0.39 | 0.133 | |
| S2 vs S4 | 0.37 | 0.063 | |
| S3vs S4 | 0.33 | 0.2 | |

**Table S6.** The relationship between environment condition and microbial community. The significance of each environment condition effect was tested with the ANOVA based on the RDA results.

| Variable | *p*-value | Raw $R^2$ | Adjusted $R^2$ |
|---|---|---|---|
| Temperature | 0.8 | 0.032 | -0.042 |
| pH | 0.003 | 0.20 | 0.139 |
| Allylthiourea | 0.002 | 0.281 | 0.226 |
| $PO_4^{3-}$-P | 0.269 | 0.09 | 0.020 |
| TN | 0.001 | 0.281 | 0.222 |
| $NH_4^+$-N | 0.009 | 0.188 | 0.12 6 |
| TOC | 0.091 | 0.128 | 0.061 |
| $PO_4^{3-}$-P Removal | 0.001 | 0.261 | 0.205 |
| TN Removal | 0.203 | 0.10 | 0.031 |
| $NH_4^+$-N Removal | 0.337 | 0.07 9 | 0.008 |
| TOC Removal | 0.321 | 0.081 | 0.010 |

**Table S7.** Mantel tests of correlations between microbial communities, environmental variables and EBPR performance. EBPR: enhanced biological phosphorus removal.

| Variable | *p*-value | *r*-value | Significance |
|---|---|---|---|
| Temperature | 0.908 | -0.22 | |
| **pH** | **0.001** | **0.51** | *** |
| **Allylthiourea** | **0.002** | **0.70** | ** |
| $PO_4^{3-}$-P | 0.700 | -0.09 | |
| **TN** | **0.001** | **0.68** | *** |
| **$NH_4^+$-N** | **0.02** | **0.38** | * |
| TOC | 0.279 | 0.09 | |
| **$PO_4^{3-}$-P Removal** | **0.001** | **0.634** | *** |
| TN Removal | 0.527 | -0.021 | |
| $NH_4^+$-N Removal | 0.388 | -0.002 | |
| TOC Removal | 0.577 | -0.076 | |

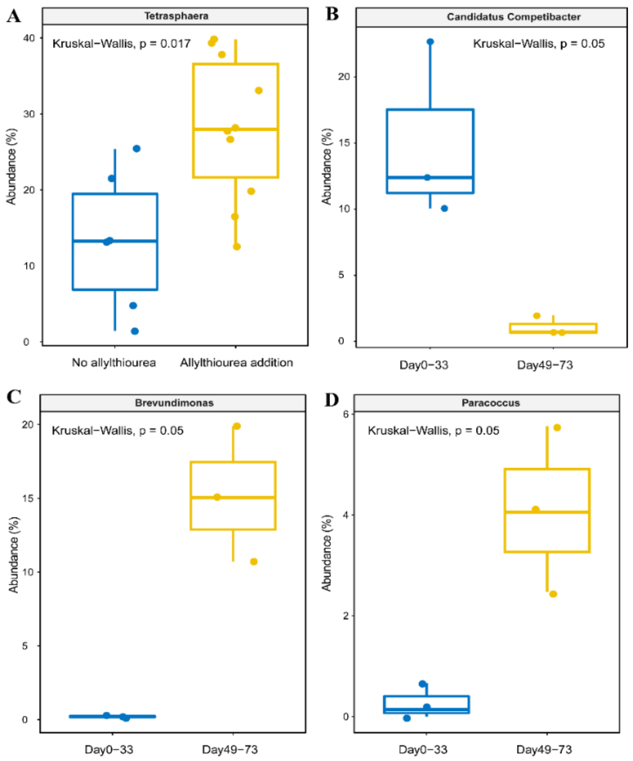


**Fig. S1** Statistics test of (A) dynamic of *Tetrasphaera* before (day 0-73) and after (day 87-157) allylthiourea addition (B) dynamic of *Candidatus* Competibacter in day 0-33 and day 49-73. (C) dynamic of *Brevundimonas* in day 0-33 and day 49-73. (C) dynamic of *Paracoccus* in day 0-33 and day 49-73.

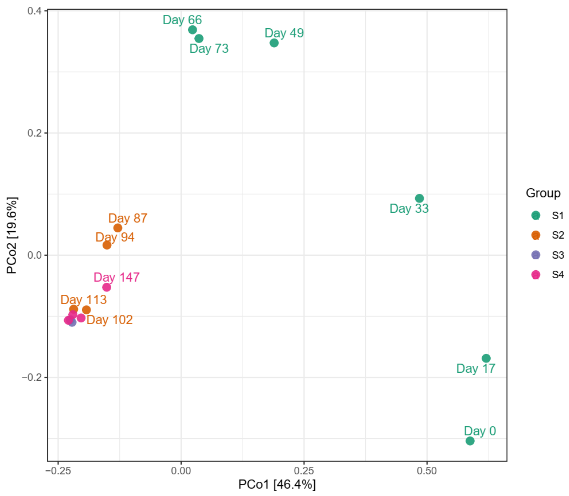


**Fig. S2** Principal coordinate analysis (PCoA) plot showing the *Tetrasphaera*-enriched microbiome establishment process. Four operational stages were defined based on the usage of nitrification inhibitor allylthiourea during the microbiome establishment process: S1: Natural enrichment phase (day 0-73) with no inhibitor; S2: Reinforced enrichment phase (day 74-120) with 1 mg/L allylthiourea as inhibitor; S3: Transient shock phase (day 121-127) with 5 mg/L allylthiourea as inhibitor); S4: Recovery phase (day 128-170) with allylthiourea dosage adjusted back to 1 mg/L.

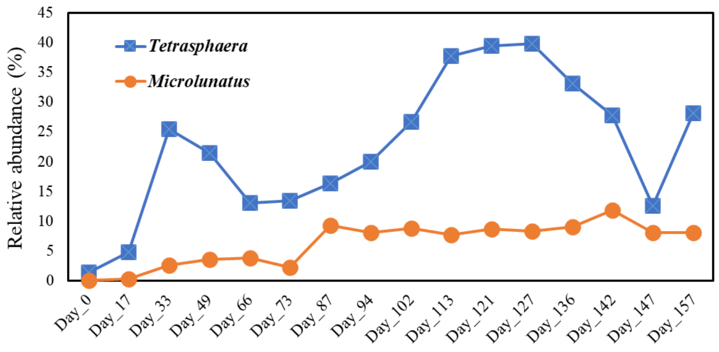


**Fig. S3** The microbial dynamic of *Tetrasphaera* PAOs and *Microlunatus* PAOs during the whole establishment process of the *Tetrasphaera*-enriched microbiome.
